# Disease diagnosis via sperm motility variations induced by body fluids in bioassay

**DOI:** 10.64898/2026.02.04.703772

**Authors:** Sihan Chen, Tiantian Liang, Mingjie Sun, Peng Wu, Yingxue Lin, Bo Yang, Chunmei Cui, Kun Lv, Chengliang Zhu, Qinghua Cui

**Author notes:** To whom the correspondence should be addressed: Bo Yang, Chunmei Cui, Kun Lv, Chengliang Zhu, Qinghua Cui. These authors contributed equally to this work (Sihan Chen, Tiantian Liang).

## Abstract

Precise diagnosis of high-risk conditions such as cardiovascular and cerebrovascular diseases still remains a challenge. We previously developed a microfluidic chip for sperm selection and observed that sperm motility is highly sensitive to environmental changes. Building on this finding, we hypothesized that motility traits of sperm could be differentially modulated by body fluids from healthy versus diseased individuals, thereby serving as potential biomarkers for disease diagnosis. To test this hypothesis, we designed a diagnostic system in which mouse sperm were co-incubated with serum samples from patients with myocardial infarction, cerebral infarction, and pancreatitis, along with matched healthy controls. Key kinematic parameters—including motility rate (MR), curvilinear velocity (VCL), straight-line velocity (VSL), linearity (LIN), and amplitude of lateral head displacement (ALH)—were analyzed using a multiparameter sperm quality analysis system. The results revealed that disease-specific serum induced distinct and reproducible changes in sperm motility patterns, enabling accurate discrimination between healthy and pathological conditions. Evaluation of these motility parameters demonstrated high diagnostic performance, with area under the receiver operating characteristic curve (AUC) values ranging from 0.719 to 0.888. This sperm-based bioassay offers a non-invasive, rapid, and cost-effective platform for disease detection and personalized health assessment, with the potential to complement existing diagnostic approaches.

## 1. Introduction

High-risk diseases such as cardiovascular and cerebrovascular diseases, malignancies, and severe inflammatory or metabolic conditions represent significant global health challenges due to their high incidence, mortality, and disability rates. Cardiovascular diseases alone account for approximately 17.9 million deaths each year, while cancers cause nearly 10 million deaths worldwide, with lung cancer being the leading oncological cause of mortality^1,2^. The enormous health and economic burdens associated with these diseases are further intensified by their frequently insidious onset and rapid progression. Many remain asymptomatic until reaching advanced stages, when treatment efficacy declines and irreversible damage has already occurred. For example, the five-year survival rate for many cancers increases from less than 20% at stage IV to more than 90% when detected at stage I^3, 4^. Similarly, timely intervention in acute cardiovascular events can reduce mortality by up to 30%^5^. These observations highlight that timely detection is the most effective strategy to improve prognosis and reduce healthcare costs^6, 7^. However, existing diagnostic technologies remain inadequate for early and accessible detection. Imaging-based methods, while indispensable, face limitations in specificity, radiation exposure, and cost. For instance, low-dose computed tomography screening for lung cancer shows false-positive rates exceeding 20% in some trials, resulting in unnecessary invasive procedures and psychological distress^8, 9^. Biochemical marker assays, such as troponin for myocardial infarction or prostate-specific antigen for prostate cancer, often lack sufficient sensitivity for early-stage detection, and their diagnostic accuracy is further compromised by cross-reactivity and physiological variability^10^. Moreover, advanced diagnostic tools are largely inaccessible in resource-limited regions, where an estimated 75% of the global population has limited access to basic radiological services. The global economic burden of cancer alone is projected to surpass 450 billion US dollars annually by 2030, much of which is attributed to delayed diagnosis and treatment. These limitations underscore the urgent need for innovative, cost-effective, and non-invasive diagnostic strategies that can identify diseases at their earliest and most treatable stages.

In recent years, cell-based biosensing systems have emerged as a promising alternative to traditional biomarker-based diagnostics. By exploiting the intrinsic sensitivity of living cells to changes in their microenvironment, these systems can detect subtle biochemical and biophysical alterations associated with pathological conditions^11^. For instance, circulating tumor cells serve as live liquid biopsies, retaining genomic and phenotypic tumor information for real-time monitoring and single-cell analysis^12^. Synthetic biology approaches further enable the engineering of cellular biosensors, such as TGF-β-responsive reporter cells that produce detectable fluorescent signals upon exposure to disease-specific molecules^13^. Notably, the indicator cell analysis platform (iCAP) utilizes cultured cells to amplify complex biomarker signals derived from serum. In a blinded validation study, the lung cancer-specific iCAP demonstrated high sensitivity and specificity in distinguishing patients with non-small cell lung cancer from those presenting with benign nodules^14^. Among various cell types, sperm cells exhibit unique and quantifiable motility characteristics, making them suitable candidates for functional biosensing^15, 16^. Our previous studies using microfluidic-based sperm sorting technology revealed that sperm motility parameters are highly responsive to minimal changes in their surrounding medium, suggesting their potential role as natural biological sensors^17^.

Building on these findings, we developed a novel diagnostic and screening system that utilizes sperm motility traits as diagnostic indicators (Figure 1). In this system, sperm cells were exposed to serum samples from patients with various diseases, including myocardial infarction (MI), cerebral infarction (CI), and pancreatitis. Kinematic responses of sperm were recorded and analyzed using advanced motion-tracking algorithms. Distinct and reproducible motility patterns were observed in response to different pathological conditions, enabling accurate disease detection through statistical modeling and machine learning. This method differs fundamentally from conventional biomarker detection by relying on functional biological responses as diagnostic readouts. It provides a non-invasive, and low-cost biosensing platform that leverages the natural sensitivity of sperm to detect disease-specific biochemical signatures.

**Figure 1:**
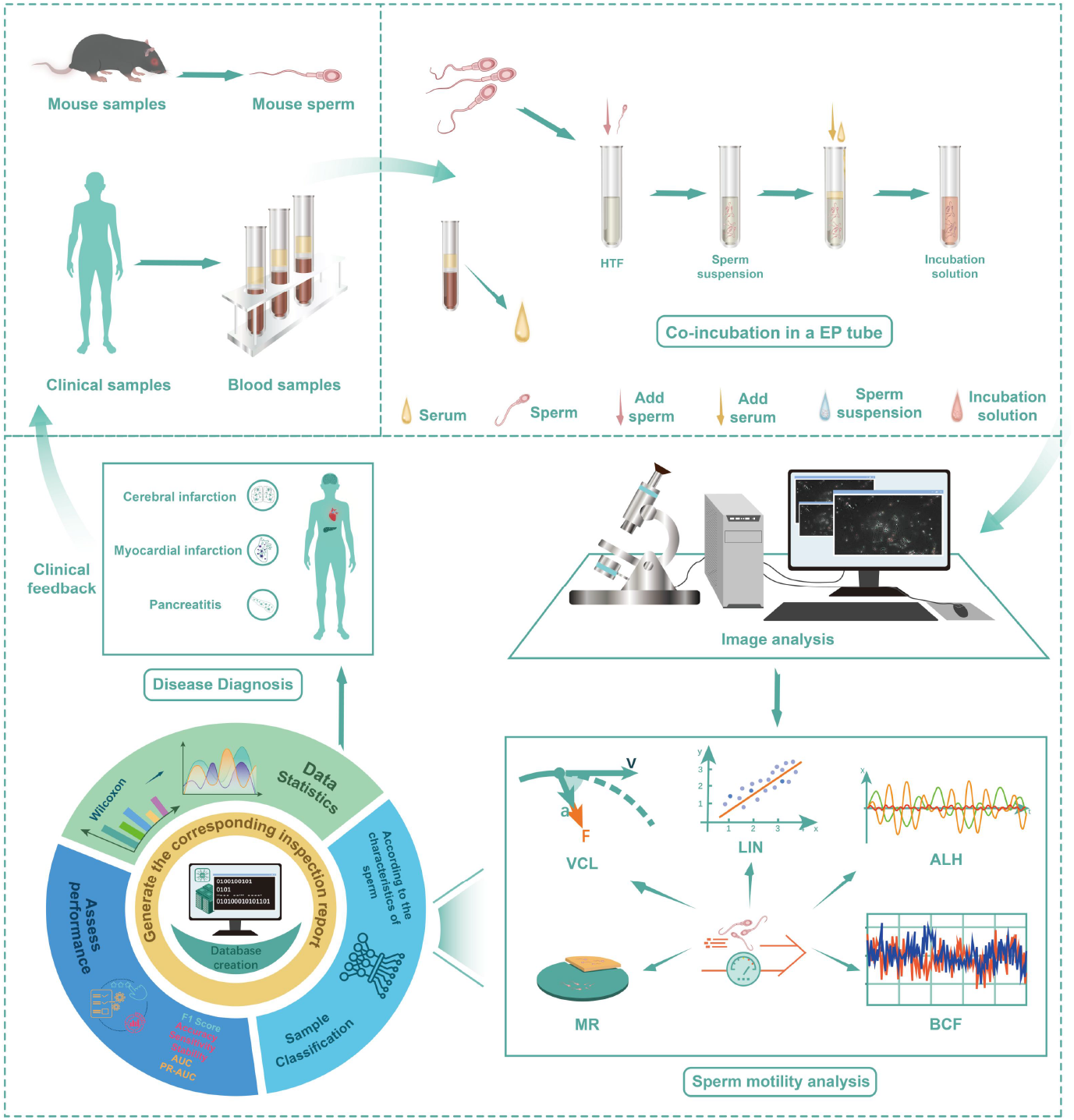
Schematic view of a novel disease diagnostic and screening system via trait variations of sperm induced by serum from pathological and healthy individuals.

## 2. Materials and methods

### 2.1 Collection of blood samples

This study was approved by the Institutional Review Board, and all participants provided written informed consent before enrollment. Serum samples were obtained from two collaborating hospitals. At the First Affiliated Hospital of Wannan Medical College, the cohort included 28 patients diagnosed with CI and 19 with MI. From Renmin Hospital of Wuhan University, sex- and age-matched control samples were collected, including 28 healthy controls for the CI group and 17 for the MI group. In addition, paired serum samples were collected from 11 patients with pancreatitis and 11 controls in this hospital.

### 2.2 Blood biochemical analysis

We first explored whether sperm motility is associated with serum biochemical parameters. To do so, we analyzed serum samples from 10 healthy individuals. Venous blood samples were collected from all these participants after an overnight fast. The collected blood samples were centrifuged to separate serum, and the subsequent biochemical analyses were performed using an automated biochemistry analyzer. A comprehensive panel of biomarkers was assessed, encompassing liver function, renal function, lipid profile, glucose metabolism, and specific proteins. The measured parameters included: alanine aminotransferase (ALT), aspartate aminotransferase (AST), alkaline phosphatase (ALP), total protein (TP), albumin (ALB), total bilirubin (TBIL), direct bilirubin (DBIL), urea, creatinine (Cr), uric acid (UA), glucose (Glu), total cholesterol (TCh), triglycerides (TG), high-density lipoprotein cholesterol (HDL-Ch), and low-density lipoprotein cholesterol (LDL-Ch). Furthermore, calculated ratios and derived indices were determined, including the AST/ALT ratio, globulin (GLB), albumin-to-globulin ratio (A/G), urea-to-creatinine ratio, total cholesterol-to-HDL-Ch ratio, as well as lipoprotein(a) Lp(a), serum sialic acid (SA), estimated glomerular filtration rate (eGFR), and adiponectin (ADPN). All procedures were strictly conducted in accordance with the manufacturer’s instructions and standard operational protocols for the analyzer and assay kits.

### 2.3 Preparation of serum samples

All blood samples were collected before any therapeutic intervention and after confirmation of the clinical diagnosis. For each participant, 5 mL of peripheral blood was drawn and allowed to stand at room temperature for 30 minutes to prevent hemolysis. Samples were then centrifuged at 4000 rpm for 5 minutes at 4°C. The resulting layers consisted of serum at the top, a gel separator in the middle, and the blood clot at the bottom. The serum layer (1 mL) was carefully aspirated into sterile, labeled tubes, sealed with parafilm, and stored at −80°C until further analysis. All sample collection and processing steps followed standardized protocols to minimize technical variation and ensure consistency and reliability for subsequent experiments.

### 2.4 Preparation of sperm samples

Sperm samples were collected from three-month-old *C57BL/6* male mice that were euthanized by cervical dislocation. The cauda epididymides were dissected and placed into pre-warmed human tubal fluid (HTF) *(Vitrolife, G-IVF PLUS/60 mL-10136)* to allow sperm to swim up. Motile sperm were collected and standardized to a concentration, then incubated in 5% CO_2_ at 37°C for 30 minutes to achieve capacitation. All procedures involving animals were conducted in accordance with institutional ethical standards and were approved by the Experimental Animal Management Committee of Wuhan Sports University (Animal Use Protocol No. 2025007). Every effort was made to minimize animal suffering and reduce the number of animals used in the study.

### 2.5 Co-Incubation of sperm suspension with serum

Before co-incubation, sperm quality was evaluated by motility assessment. Samples exhibiting approximately 40% motility and a concentration of 1 × 10^6^ sperm per mL were selected for subsequent experiments. The sperm suspension was then mixed with serum at a 10:1 volumetric ratio in sterile Eppendorf tubes and incubated for 10 minutes in a humidified incubator at 37°C with 5% CO_2_. This standardized procedure ensured controlled biochemical interaction under physiological conditions and maintained experimental consistency and reproducibility.

### 2.6 Analysis of sperm motility

Sperm motility was analyzed using a clinical-grade multiparameter sperm quality analysis system *(BEION, V4*.*20)*, which employs advanced image recognition and particle-tracking algorithms to identify and trace sperm head movements frame by frame within the microscopic field. This system allows precise quantification of detailed kinematic parameters, including motility rate (MR), which represents the percentage of motile sperm and reflects overall vitality; curvilinear velocity (VCL), defined as the average velocity of the sperm head along its actual curved trajectory; straight-line velocity (VSL), referring to the velocity along a direct line from the starting to the ending point; and average path velocity (VAP), representing the velocity along the time-averaged trajectory. Linearity (LIN), calculated as the ratio of VSL to VCL, indicates the straightness of the trajectory. In contrast, straightness (STR), the ratio of VSL to VAP, reflects the efficiency of forward movement. Wobble (WOB), determined by the ratio of VAP to VCL, describes the oscillation amplitude of the sperm head around its mean path, and the amplitude of lateral head displacement (ALH) quantifies the maximum lateral deviation of the sperm head from that path. Beat-cross frequency (BCF) measures the number of times the sperm head crosses its average trajectory per unit time, indicating the frequency of flagellar beats. Finally, mean angular displacement (MAD) corresponds to the time-averaged absolute turning angle of the sperm head, with higher values indicating greater instability in the direction of movement.

### 2.7 Statistical analysis and performance evaluation

Comprehensive information was recorded in Supplementary Table S1, including hospital of origin, patient’ sex and age, disease type, and post-incubation sperm motility parameters. Each sample was assigned a unique identifier to ensure traceability and prevent data duplication. Quality control checks were implemented to validate parameter ranges and unit consistency. Wilcoxon test was employed to compare post-incubation sperm motility parameters between the serum samples of patients and those from healthy control groups, with significance levels set at *p < 0.05, **p < 0.01, ***p < 0.001, ****p < 0.0001. To evaluate the diagnostic performance of the proposed system, several statistical indicators were used, including the area under the receiver operating characteristic curve (AUC), the area under the precision-recall curve (PR-AUC), sensitivity, precision, accuracy, and F1-score. Figures were generated using the Seaborn library in Python, and statistical analyses were performed using the stats module^18^.

## 3. Results

### 3.1 Serum-induced alterations in sperm motility

Serum samples were collected from patients and age- and sex-matched healthy controls, including 28 cases of CI and 28 controls, 19 MI patients and 17 controls, and 11 pancreatitis patients and 11 controls. The mean age across all groups was over 60 years, with demographic characteristics provided in Table 1. Each serum sample was independently introduced into the diagnostic system to evaluate alterations in sperm motility characteristics. Representative motion trajectories of sperm following exposure to various serum types are shown in Figure 2. Correlation analysis revealed that serum Lp(a) levels showed strong positive associations with sperm motility metrics, with the highest correlation observed for VSL (Rho = 0.93, P-value = 1.3e-4), followed by VAP (Rho = 0.89, P-value = 1.38e-3), VCL (Rho = 0.79, P-value = 9.84e-3), and ALH (Rho = 0.88, P-value = 1.98e-3) (Figure 3), indicating that sperm motility is responsive to serum biochemical composition. Comparative analysis further showed that MR differed significantly between controls and patients in both the CI group (mean value: 23.98% for controls vs. 10.04% for disease, P-value = 6.04e-7, Wilcoxon’s test) and MI group (mean value: 17.66% vs. 8.48%, P-value = 3.64e-4, Wilcoxon’s test), whereas the pancreatitis group exhibited a weak yet non-significant decreasing trend (22.43% vs 17.76%, P = 0.082) (Figure 4A). These findings suggest the potential utility of sperm motility analysis in discriminating between patients with complex diseases and healthy individuals.

**Table 1.** Basic information of samples.

| Disease | Type | No. of sample (% male) | Mean of Age | Std of Age |
| --- | --- | --- | --- | --- |
| CI | Control | 28 (71.43%) | 67.96 | 10.75 |
|  | Disease | 28 (71.43%) | 68.29 | 11.23 |
| MI | Control | 19 (78.95%) | 62.42 | 12.17 |
|  | Disease | 17 (82.35%) | 61.88 | 13.00 |
| Pancreatitis | Control | 11 (72.73%) | 64.55 | 8.63 |
|  | Disease | 11 (72.73%) | 64.64 | 8.57 |

**Figure 2:**
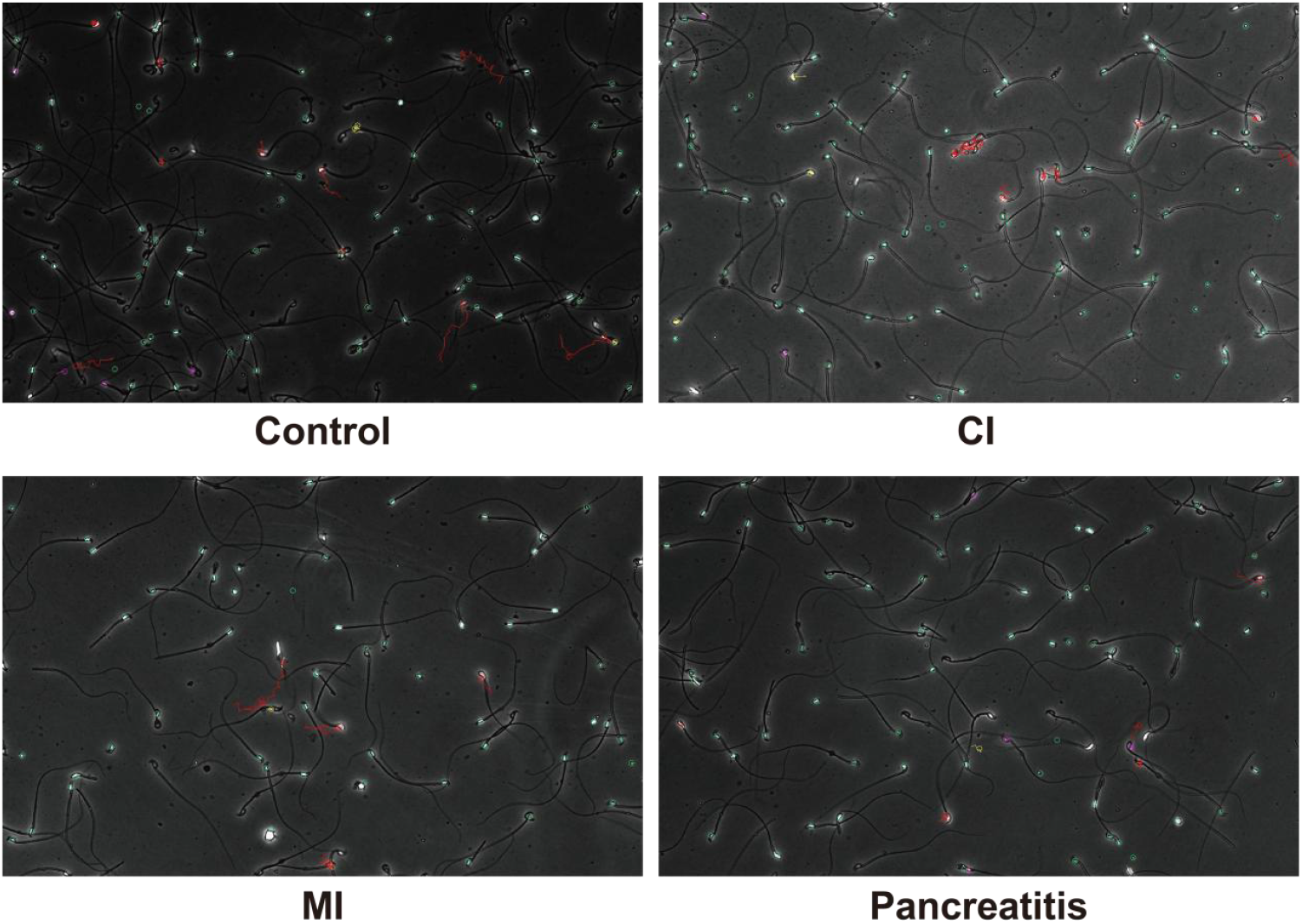
Representative images of sperm movement trajectories following serum intervention from diverse origins.

**Figure 3:**
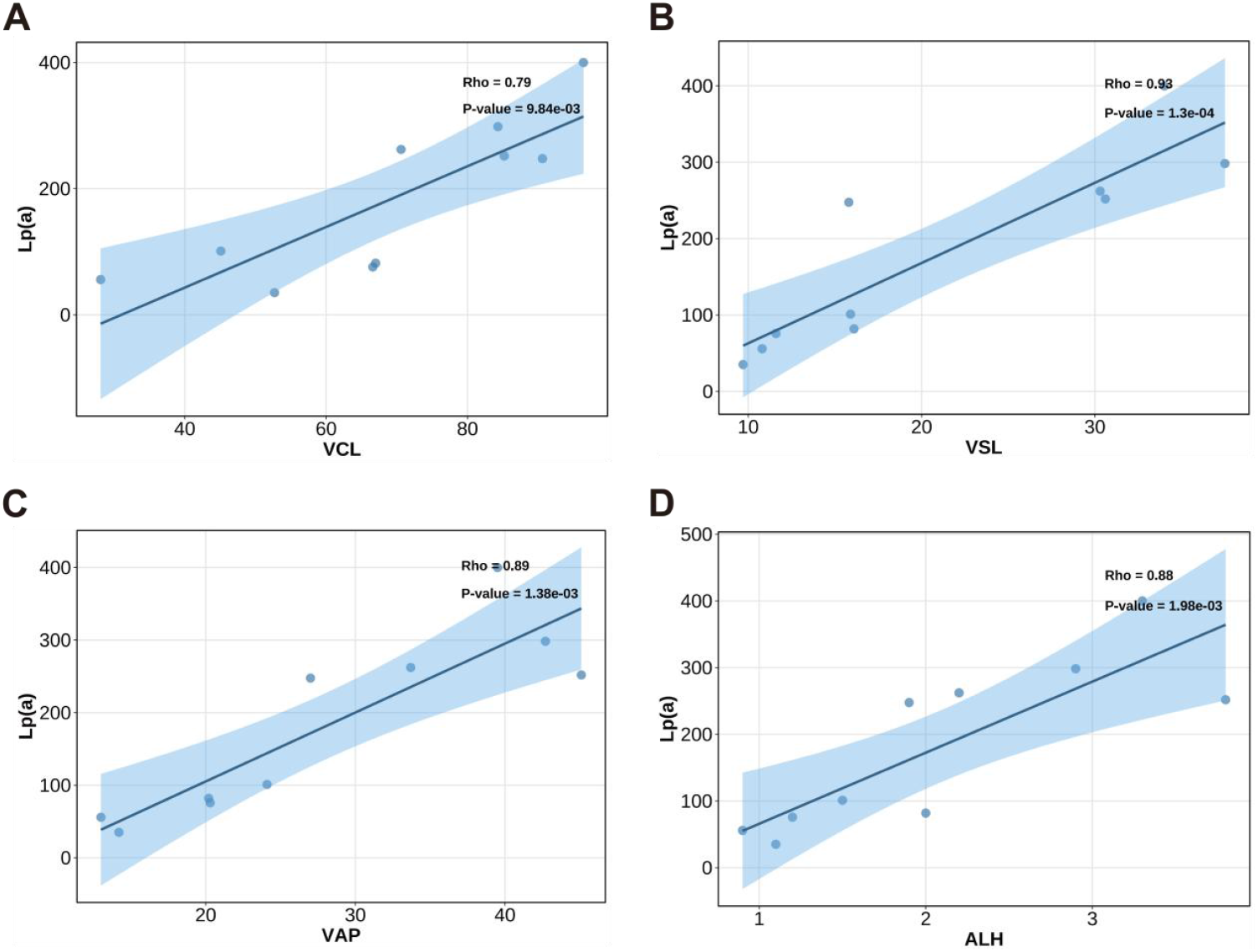
Correlation analysis between sperm motility parameters and serum biochemical indicators. The motility parameters and serum biochemical levels were measured and compared across four subgroups divided based on serum lipoprotein(a) concentrations. (A) Curvilinear velocity. (B) Straight-line velocity. (C) Average path velocity. (D) The amplitude of lateral head displacement.

**Figure 4.**
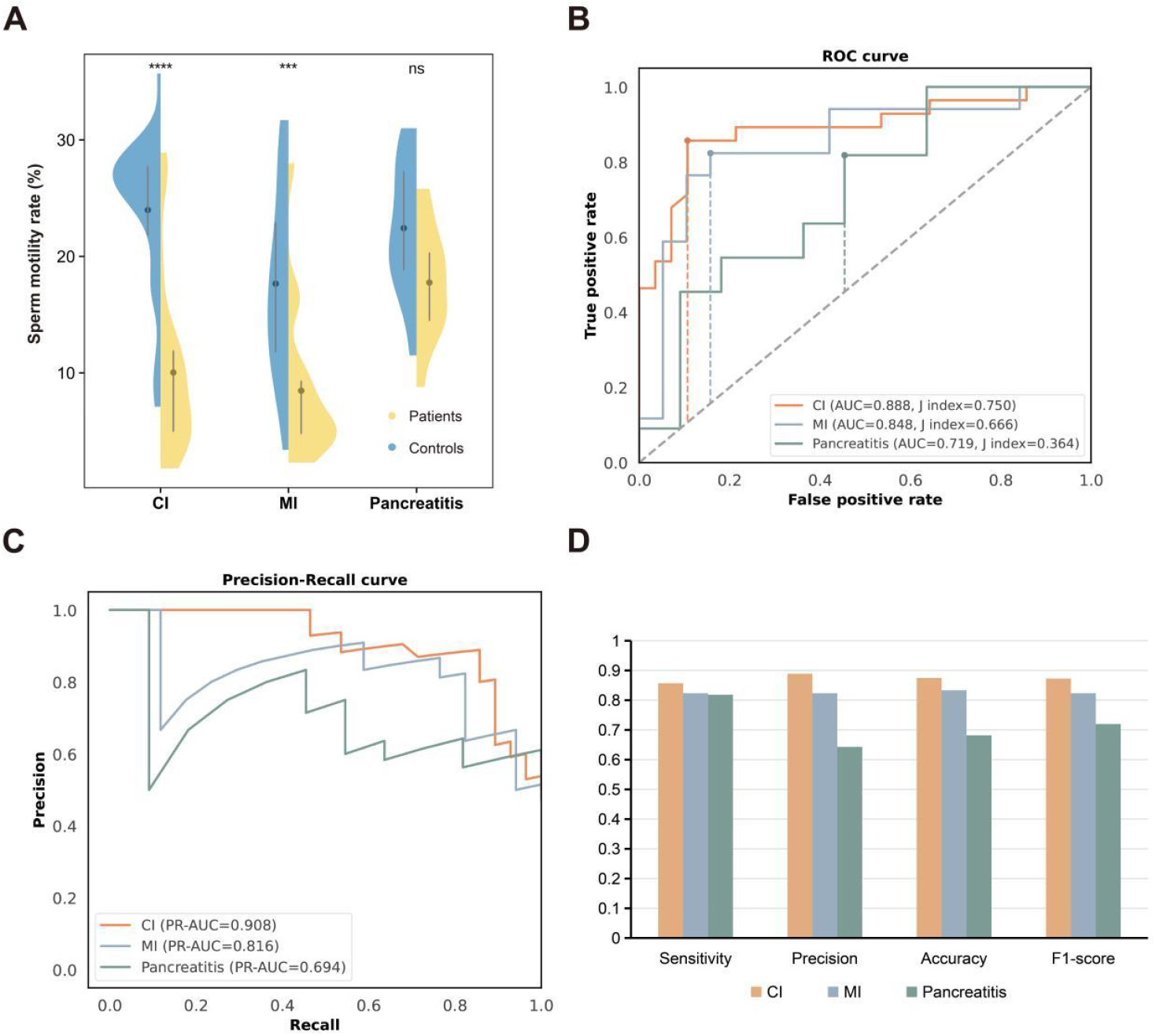
Diagnostic performance of sperm motility rate (MR) in disease classification. (A) MR distributions across patient groups and controls. (B) ROC curves for CI, MI, and pancreatitis classification. (C) Precision-recall curves for each disease group. (D) Main classification metrics at optimal MR thresholds.

### 3.2 Diagnostic performance evaluation

The diagnostic performance of sperm motility parameters was further evaluated using receiver operating characteristic (ROC) and precision-recall (PR) analyses. Notably, sperm MR demonstrated excellent classification capability, with AUC values of 0.888 for CI, 0.848 for MI, and 0.719 for pancreatitis, alongside PR-AUC values of 0.908, 0.816, and 0.694, respectively (Figure 4B and 4C). Meanwhile, optimal classification thresholds of MR determined by Youden’s index yielded robust performance metrics: for CI, MI, and pancreatitis, sensitivity was 85.7%, 82.4%, and 81.8%; precision was 88.9%, 82.4%, and 64.3%; accuracy was 87.5%, 83.3%, and 68.2%; and F1-scores were 0.873, 0.824, and 0.720 (Figure 4D), respectively. Additionally, other parameters, such as VCL and MAD, showed slightly inferior classification ability (Supplementary Table S2). Together, the system demonstrated superior diagnostic performance across diverse disease conditions, highlighting its efficacy in identifying diseased individuals.

## 4. Discussion

This study established a novel diagnostic system that uses sperm motility characteristics as dynamic biomarkers to identify individuals at high risk of disease. The results clearly demonstrated that serum exposure from patients with various pathological conditions markedly alters sperm kinematic behavior. These changes enabled accurate differentiation between healthy individuals and patients with CI, MI, and pancreatiti with AUC values ranging from 0.719 to 0.888, confirming the system’s robust diagnostic capability. The observed alterations in motility likely result from complex pathophysiological changes in serum composition. Factors such as metabolites, inflammatory cytokines, lipids, and extracellular vesicles may influence sperm activity by interacting with surface receptors or regulating intracellular signaling cascades that control flagellar motion^19, 20^. This mechanism highlights the sensitivity of sperm motility as an integrative biological sensor that reflects systemic physiological disturbances. In contrast to conventional biomarker detection strategies that rely on single molecular targets, this approach captures comprehensive cellular responses, providing a functional and holistic view of disease-related biochemical changes^21,22^. The proposed platform thus represents a promising and potential alternative for disease screening and monitoring. It offers advantages including rapid analysis, minimal sample processing, and low instrumentation requirements, which make it particularly suitable for large-scale population screening and applications in resource-limited settings. Moreover, the use of living cells as active biosensors introduces a new dimension to precision diagnostics, bridging the gap between molecular information and functional physiological outcomes. Future investigations should broaden the clinical and biological scope of this system. Firstly, the mechanisms why this system works remain unknown and should be explored. Further integration with proteomic and metabolomic analyses could clarify the molecular mechanisms linking serum components to alterations in sperm motility^23, 24^. Secondly, evaluating its diagnostic performance across a broader range of diseases—including cancer, autoimmune, neurodegenerative, and rare genetic disorders—will help establish its general applicability. In addition, it is also necessary to test this system on other diagnosis tasks such as disease subtyping, risk prediction, classification, stratification, phenotyping, drug response prediction, and prognosis. Another important issue is to test other body biofluids, such as plasma, urine, tear, sweat, saliva, or fecal extracts, may enhance accessibility and enable non-invasive disease detection, although compositional variability will necessitate optimized preprocessing and standardized analytical frameworks. Technological innovation will also be crucial for system refinement. The development of automated microfluidic platforms integrating sample loading, motility tracking, and data processing could substantially improve throughput and reproducibility. Advanced computational tools, particularly deep learning algorithms for motion pattern recognition, may further enhance classification accuracy and robustness^25, 26^. To validate the consistency of performance across diverse populations and clinical settings, multi-center clinical studies are needed to validate performance consistency across diverse populations and clinical settings, ensuring reproducibility, minimizing operator-dependent variability, and establishing reliable diagnostic criteria^27^. Looking forward, it is worthwhile to explore alternative biosensing entities beyond mouse-derived sperm. Promising candidates include embryonic stem cell-derived germ cells, as well as organisms such as *C*.*elegans* and microorganisms (e.g. *E*.*coli* and *Plasmodium*).

In summary, this sperm-based diagnostic system offers an innovative, biologically inspired approach to disease detection. By transforming cellular motility behavior into measurable diagnostic signals, it provides a powerful and accessible approach for early identification of pathological conditions. Continued interdisciplinary efforts combining biological insight, technological development, and clinical validation will be essential to unlock its full potential for personalized and preventive healthcare.

## 5. Conclusions

This study established and validated a novel disease diagnostic and screening system that utilizes sperm motility characteristics as sensitive functional biomarkers. The findings demonstrate that alterations in sperm kinematic behavior induced by patient serum can accurately distinguish various disease conditions, highlighting the system’s capability and potential for early and reliable detection. By transforming cellular motion into quantifiable diagnostic information, this approach provides a biologically integrated and non-invasive method for assessing systemic health conditions.

## Ethical approval

The study was approved by the Medical Ethics Committee of Renmin Hospital of Wuhan University (IRB Reference Number: WDRY2025-K212) and Yijishan Hospital (IRB Reference Number: 2013110017).

## Data availability

All data supporting the findings of this study are included in the article and supplementary materials and can be obtained from the corresponding author upon request. Source data are provided with this article.

## Acknowledgements

This study was supported by the grants from the National Natural Science Foundation of China [82470373, 32301239, 62025102] and Natural Science Foundation of Anhui Province [2408085JX011]. We thank Profs. Yu Zhang and Yang Li at Peking University for valuable discussion on embryonic stem cell-derived germ cells.

## Authors’ contributions

Q.C. proposed the idea. S.C., T.L, M.S., performed the experiment. P.W. processed the serum samples. C.C. analyzed the data. B.Y., C.C., K.L., C.Z., and Q.C. supervised the study. S.C. wrote the draft manuscript. Q.C. edited the manuscript. All authors read and approved the final manuscript.

